# Virus-mediated recycling of chemoautotrophic biomass

**DOI:** 10.1101/2025.05.27.656380

**Authors:** Elaine Luo, Ngoc D. Pham, Timothy J. Rogers, Joseph J. Vallino, Bayleigh E. Benner, Gareth Trubl, Julie A. Huber

## Abstract

Aquatic environments absorb ∼2.5 gigatonnes of atmospheric carbon each year^1^, more than the carbon stored in the atmosphere, soils, and all biomass combined. Primary producers transform this dissolved inorganic carbon into biomass that can subsequently flow into other trophic levels, or be released back into the environment through viral lysis. While there is substantial knowledge about the diversity and activity of viruses infecting photoautotrophic primary producers, little is known about viruses infecting chemoautotrophs, representing a gap in our understanding of key microbial processes driving global carbon cycles. Here, we combine metagenomics with ^12/13^C stable isotopic probing mesocosm experiments in a marine-derived meromictic pond to quantify lineage-specific carbon cycling activity to identify key microbial populations driving carbon cycling. We then tracked the flow of carbon from active chemoautotrophs to their viruses and found evidence supporting virus-mediated recycling of chemoautotrophic biomass through the production of viral particles. In particular, active populations of hydrogen/sulfur-oxidizing chemoautotrophs (*Thiomicrorhabdus, Hydrogenovibrio, Sulfurimonas, Sulfurovum*) were targeted by viruses. Considering the widespread distribution of chemoautotrophs on Earth, we postulate that this previously overlooked component of the microbial carbon cycle is a globally relevant process that has implications for our planet’s carbon cycle. This work provides the foundation for revealing the role of viral lysis in chemoautotrophic primary production and builds toward biogeochemical models that incorporate viral recycling of chemoautotrophic biomass.

**Summary statement:** The diversity, mechanisms, and processes governing microbial primary production and the recycling of autotrophic biomass are fundamental to our planet’s carbon cycle. These processes have implications for carbon sequestration, ocean biogeochemistry, and the overall balance of carbon dioxide in the atmosphere. Beneath the Earth’s sunlit layer, primary production is driven by microbial chemoautotrophs that derive energy from the oxidation of reduced compounds, such as hydrogen and sulfur, to form the base of the food web. Growing evidence suggests that aquatic ecosystems fueled by chemoautotrophy are widely distributed on Earth, ranging from beneath ice shelves to coastal upwelling regions to oxygen minimum zones, deep-sea hydrothermal vents and cold seeps, groundwater, and meromictic ponds and lakes^2–7^. Studying microbial processes regulating chemoautotrophic primary production and the recycling of chemoautotrophic biomass is fundamental to our understanding of global carbon cycles.

While the diversity, function, and activity of viruses targeting photoautotrophs have been well-described across aquatic ecosystems^8^, we have little understanding of viruses involved in the recycling of chemoautotrophic biomass. Viruses are a major source of cellular mortality and carbon cycling in aquatic environments^9–11^. Viral lysis is estimated to transform ∼150 gigatonnes of carbon annually from biomass back into the environment, equivalent to ∼25 times that of the ocean’s biological carbon pump^12,13^. Despite recognition of the important role of viruses in aquatic habitats, there is a large gap in our understanding of the impact of viruses on globally distributed chemoautotrophs^2,4,5,14–18^. In this study, we show that viruses are not merely passive players but active agents recycling carbon fixed by productive chemoautotrophs, fundamentally reshaping how we view carbon and nutrient cycling in redox-active ecosystems.

## Quantifying population-specific carbon cycling activity

Although metagenomic studies have the potential to discover novel microbial diversity, linking viruses to their biogeochemical impacts remains an ongoing challenge in environmental microbiology. Metagenomic studies have identified putative viruses infecting chemoautotrophic hosts across oxygen minimum zones, mesopelagic open ocean, and deep-sea hydrothermal vents^19–26^. Lysogeny was hypothesized to be the dominant mode of infection, based on metagenomic predictions on microbial communities at deep-sea hydrothermal vents and in the mesopelagic ocean^19–22^. These *in silico* predictions, however, cannot definitively link a virus to its host, cannot determine whether a virus or host is active, and cannot identify whether this interaction is ecologically or biogeochemically relevant. Whether and which viruses play a role in the active recycling of chemoautotrophic biomass in aquatic environments remains to be determined. These challenges highlight the need to develop new experimental approaches to provide a mechanistic understanding of the role of viruses in microbial carbon cycling.

Here, we combine qualitative stable isotope probing (qSIP) and metagenomics to track the flow of carbon across natural microbial populations. Combining SIP with metagenomics provides critical mechanistic links between novel genomic diversity, function, and activity in ecosystem processes^27–30^. Since viruses mostly use host nucleotides for their genomes during replication^31^, we expect tight correlations in isotopic signatures between viral and host genomes^29^, enabling a reference-independent approach to link novel viruses to their hosts. We conducted ^12/13^C SIP mesocosm experiments using dissolved inorganic carbon (DIC) and samples collected from the chemocline of a marine-derived meromictic pond as a model ecosystem for chemoautotrophic communities (Fig. S1, S2). We quantified the carbon cycling activity of microbial populations and identified key chemoautotrophs and viruses responsible for cycling carbon in this system. Through the simultaneous collection of both cellular (>0.22μm) and virus-enriched (0.02-0.22μm) samples, we found that viruses infect productive chemoautotrophs and actively cycle carbon through cell lysis and the production of new viral particles.

Prior SIP-metagenomic studies have generally sequenced only the light and heavy fractions of DNA density gradients between treatments, which enables a binary distinction between organisms that have and have not taken up the labeled substrate with a predicted threshold. Sequencing multiple density fractions, on the other hand, can enable the quantification of carbon cycling activity for each microbial population^29^. Here, we sequenced five density fractions (Fig. S3) to enable the quantification of isotopic enrichment through the calculation of ^13^C Excess Atom Fraction (EAF) for each cellular and viral population genome. We also included a ^12^C control, which accounts for baseline variability in genomic density (*e.g.*, based on GC content), to calculate the difference in density observed between the ^12^C (^13^C natural abundance) and ^13^C enriched treatments, and enable quantitative measurements of isotopic enrichment as a proxy for carbon cycling activity for each microbial population.

## Key microbial drivers of chemoautotrophic primary production

Our results indicate that under experimental conditions, dark carbon fixation in this marine-derived meromictic coastal pond is driven by a few highly active microbial genera via sulfur and/or hydrogen oxidation. In the experiment, we determined the ^13^C isotopic enrichment of >0.45μm particulate organic carbon using gas chromatography mass spectrometry, showing that it increased from day 0 to 7 of incubation (Fig. S4). We quantified population-specific carbon cycling activity using EAF values for 28,788 cellular contigs and 187 high-confidence viral population genomes, serving as a quantitative metric of each population’s carbon cycling activity (Tables S1, S2). Significant carbon cycling was detected in 3193 cellular contigs and 35 high-confidence viral population genomes (EAF >0.049, Fig. 1). While most microbial populations demonstrated insignificant carbon cycling activity, a small subset of microbial populations were highly active (Fig. 1). The most active primary producers were sulfur and/or hydrogen-oxidizing bacteria: *Thiomicrorhabdus* and *Hydrogenovibrio* (phylum: Pseudomonadota), and *Sulfurimonas* and *Sulfurovum* (phylum: Campylobacterota). Given that species richness was comparable pre-and post-incubation (Table S3), we hypothesize that these experimental results were generalizable to the sampled environment. The relative abundances of these four highly active groups combined accounted for a small fraction (0.13%) of the total prokaryotic assemblages recovered from this environment, suggesting that these rare but key taxa would have been overlooked based on metagenomic sequencing alone.

**Figure 1.**
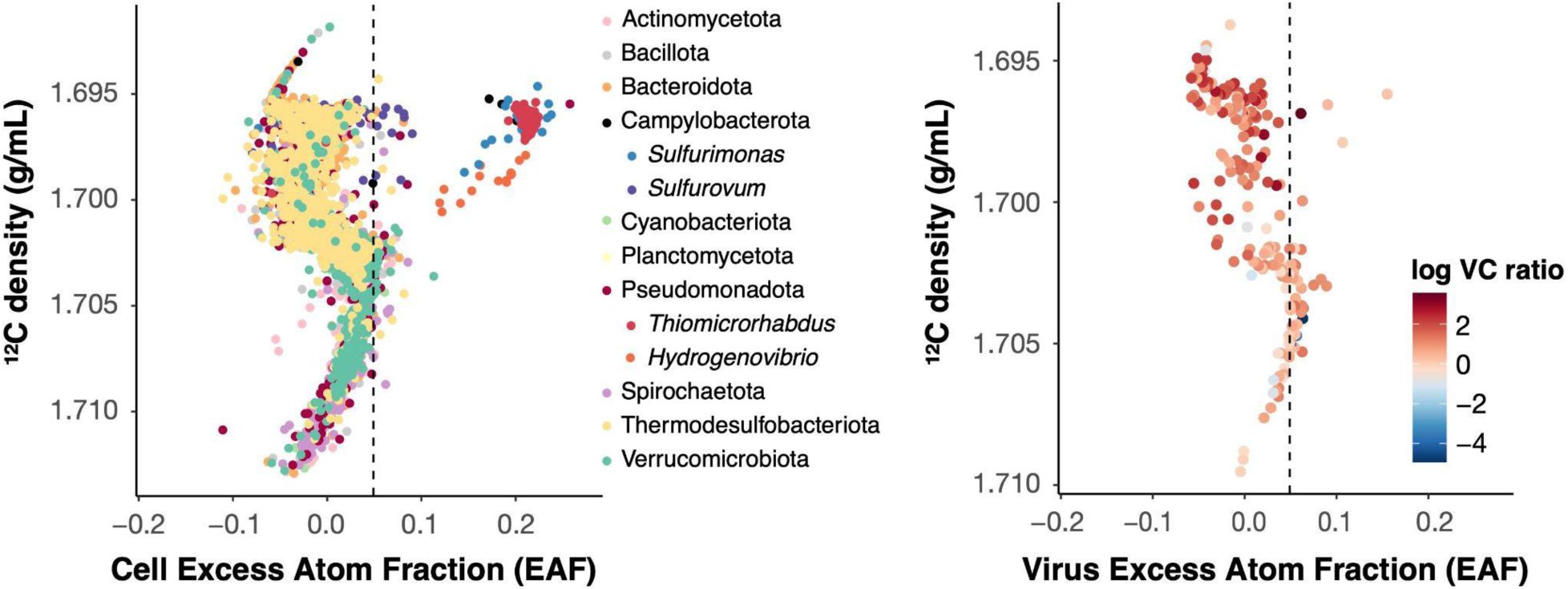
Carbon cycling activity as determined by ^13^C Excess Atom Fraction (EAF), plotted relative to the contigs’ average density in the ^12^C control, of cellular contigs from metagenome-assembled genomes (MAGs, left) and viral population genomes (right). Cellular contigs are colored by their MAG phylum-or genus-(italic) level assignments from GTDB, and only phyla with >100 contigs are visualized. Contigs above the dashed line indicated significant carbon incorporation (EAF>0.049). Viral populations are colored by the log-ratio of their relative abundance in the virus-enriched to cell-enriched sample (VC ratio) in the environmental sample, approximating their reproductive strategy. Prophages are expected to have near-zero extracellular presence (negative log VC ratio), while populations producing viral particles are expected to have detectable extracellular presence (higher log VC ratio).

Although *Thiomicrorhabdus* and *Sufurimonas* have been previously reported in the water column of meromictic ecosystems and hypothesized to contribute to sulfur cycling and carbon fixation^32–34,16^, *Hydrogenovibrio* and *Sulfurovum* have not. Here, we show that they are not only present, but actively fixing carbon. Differences in primary production were observed even within taxonomically closely-related populations within the same genus (Fig. S5). For example, 96%, 30%, 60%, and 9% of contigs identified as *Thiomicrorhabdus*, *Hydrogenovibrio*, *Sulfurimonas*, and *Sulfurovum,* respectively, were highly active (EAF >=0.2); while 0.6%, 0%, 24%, and 81%, respectively, did not show significant label incorporation (EAF <=0.049). These results indicate that primary production in this system is driven by a small subset of highly active chemoautotrophic populations that serve as keystone species for carbon cycling.

The most abundant prokaryotic populations showed insignificant carbon cycling activity (Fig. 2). Contigs assigned to the genus *Chlorobium*, *Synechecoccus*, *Desulfosarcina*, *Desulfobacter*, and *Desulfonema* respectively accounted for 4.5%, 4.0%, 3.4%, 1.1%, and 1.1% of the cellular assemblages recovered from the environmental sample. Although both *Chlorobium* and *Synechecoccus* are common primary producers across diverse aquatic ecosystems, respectively dominating anoxic and oxic depths of meromictic ecosystems^34–41^, both showed undetectable carbon fixation in the dark experimental conditions (Fig. 2). We do not expect that these photoautotrophs were active in the environment sampled, given that photosynthetically active radiation (PAR) at the time and depth of sampling, on a sunny day, was near-zero (0.04μmol/m²/s). Even extremely low-light adapted *Chlorobium* strains isolated from the Black Sea could not grow at PAR levels of 0.25μmol/m²/s ^42^. Although the lowest reported minimum light requirements for phototrophic carbon fixation was 0.015μmol/m²/s in a putative *Chlorobiacaea*, its rate of carbon fixation at these light levels estimated its doubling time at 26 years^43^. Furthermore, 0.04μmol/m²/s is 140 times lower than the lowest reported minimum light requirements for net *Synechococcus* photosynthesis^44^. Taken together, we reason that the two most dominant autotrophs may have been vertically transported from upper layers and were irrelevant to primary production at the time and depth of sampling, a conclusion supported by their lack of activity as measured by EAF. Our results show that the prevalence and predicted function of taxonomic groups, as recovered by metagenomic sequencing, do not reflect their activity in the habitat sampled. This finding has broad implications for environmental studies that rely on metagenomic sequencing alone to identify microbial function and predict their ecological and biogeochemical impacts.

**Figure 2.**
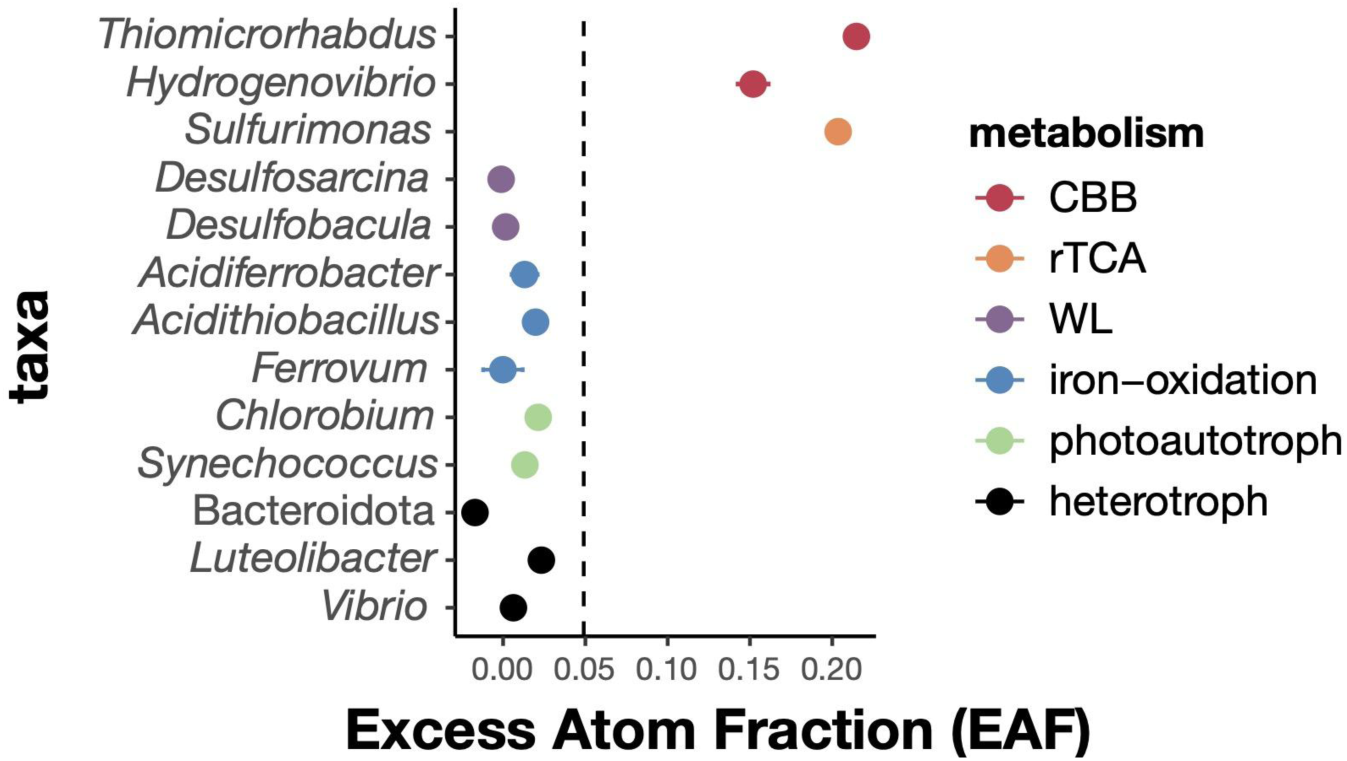
Carbon cycling activity as determined by the ^13^C Excess Atom Fraction (EAF) of MAGs encoding the following carbon fixation pathways: Calvin-Benson-Bassham cycle (CBB), reductive tricarboxylic acid cycle (rTCA), and Wood-Ljungdahl (WL). The carbon cycling activity of contigs from iron-oxidizing chemoautotrophs, photoautotrophs, and heterotrophs are shown below. The circle represents the mean EAF of all contigs in that MAG or taxa and the line represents the standard error, both color-coded by its metabolism. Genus-level taxonomic assignments are shown in italics, while the phylum-level taxonomic assignment is not. The dashed line represents significant label incorporation (EAF>0.049).

## Current limitations on linking viruses to active chemoautotrophs using *in silico* predictions

*In silico* predictions of viral taxonomy and function yielded limited information. Taxonomic identification for novel environmental viruses is hindered by the sparse representation of uncultivated viruses in reference databases. 31,448 non-redundant low-quality to complete viral populations were recovered with a sequence length of ≥5 kbp. Using a marker-protein based annotation program, the majority (92%) of viral population genomes received only class-level annotations (Fig. S6) and were novel at the order level and beyond. None received a genus-level annotation. Using protein sequence similarity searches against reference databases, only 40 (0.1%) of viral population genomes received a genus-level annotation, all broadly related to *Ostreococcus* and *Micromonas* viruses. 277 (0.9%) of total viral populations were identified as temperate phages via marker genes and alignments to putative prophages in contigs from cell-enriched samples. Using metagenomic virus-host linkage methods, such as alignments to prophages in cellular assemblies and CRISPR spacers, only 0.9% (294) were linked to cellular contigs (Fig. S6), and 205 of these yielded taxonomic annotations. Taxonomic identification of linked host contigs revealed that these viruses potentially target bacteria in the genus *Methylococcus*, *Spirochaeta*, *Desulfuromonas*, *Desulfobacter*, and *Chlorobium*, ordered from high to low EAF of viral population genomes. *In silico* host predictions did not link any viruses to active primary producers in this environment (EAF >0.049). These results highlight the current limitations of *in silico* approaches in identifying the taxonomy and function of novel viral diversity, which remains a major challenge in the field of environmental microbiology.

## Identifying viruses infecting active chemoautotrophs using their isotopic signatures

We show that similarities in isotopic signatures can be utilized to overcome the above limitations as a reference-independent method to link active viral populations to their hosts. Virus-host EAFs exhibited a strong linear correlation (N=15, P-value = 1.55e-7), as demonstrated by high-confidence viral population genomes that yielded host predictions through prophage and CRISPR spacer mapping (Fig. S7). This pattern reflects expectations that viruses mostly incorporate host nucleotides during replication. Based on this expectation, we postulate that viral populations with high carbon cycling activity in the cell-enriched size fraction (Fig. 1), and in particular those with the highest EAF values, targeted active chemoautotrophs in this environment (*Hydrogenovibrio*, *Thiomicrorhabdus*, *Sulfurimonas*, *Sulfurovum*). Viral populations with significant carbon cycling activity in the cell-enriched size fraction were rare (Fig. S7), accounting for 1.6% of the total viral assemblage recovered from the environmental sample, consistent with the expectation that they were rare but productive as they targeted highly active chemoautotrophic hosts.

## Viruses infecting productive chemoautotrophs actively cycle carbon through the production of viral particles

The simultaneous sampling of cell-enriched (>0.22μm) and virus-enriched (0.02 - 0.22μm) size fractions indicated that 99.8% of viral population genomes produced viral particles *in situ*, as demonstrated by their presence (nonzero interquartile coverage) in the virus-enriched size fraction in the environmental sample. Viral populations producing viral particles are expected to be observed in both size fractions, whereas passive prophages integrated into cellular genomes are expected to be restricted to the cell-enriched size fraction. We used the VC ratio, representing the ratio of extracellular to intracellular sequence abundance for each viral population, to show that almost all viral populations were present in the viral-particle fraction (Fig. 1). Viral particles experience decay and estimated rates of turnover range from 0.036 - 30 days across aquatic environments, averaging between 1.6 - 6.1 days^45^. As a result, we expect that viral particles sampled in the virus-enriched size fractions represent viral populations that have recently produced viral particles.

All 35 high-confidence viral populations with significant carbon cycling activity (EAF>0.049) in the cell-enriched size fraction were observed in the virus-enriched size fraction, indicating that viruses targeting hosts with high carbon cycling activity were actively cycling carbon through viral particle production. Furthermore, the isotopic enrichment of viral genomes was observed both inside cells and in viral particles, as high-confidence viral populations showed significant carbon cycling activity (EAF>0.049) in both size fractions (Fig. S7). These results reveal multiple lines of evidence showing that viral populations actively recycled chemoautotrophic biomass through the production of new viral particles, positioning them as key contributors to carbon cycling in these environments.

## Estimating population-specific viral turnover rates

The isotopic enrichment data of viral populations can enable estimates of turnover rates at the population and the community level. We expect that if a viral population completely turned over in the viral particle pool during our incubation period, its isotopic enrichment in the virus-enriched sample would reflect its isotopic enrichment in the cell-enriched sample (i.e. 1:1 ratio). However, the EAF values of viral populations in the virus-enriched samples were generally below 1:1 compared with their EAF values in the cell-enriched samples (Fig. 3), indicating that the standing stock of viral particles did not completely turn over during the 7-day incubation period.

**Figure 3.**
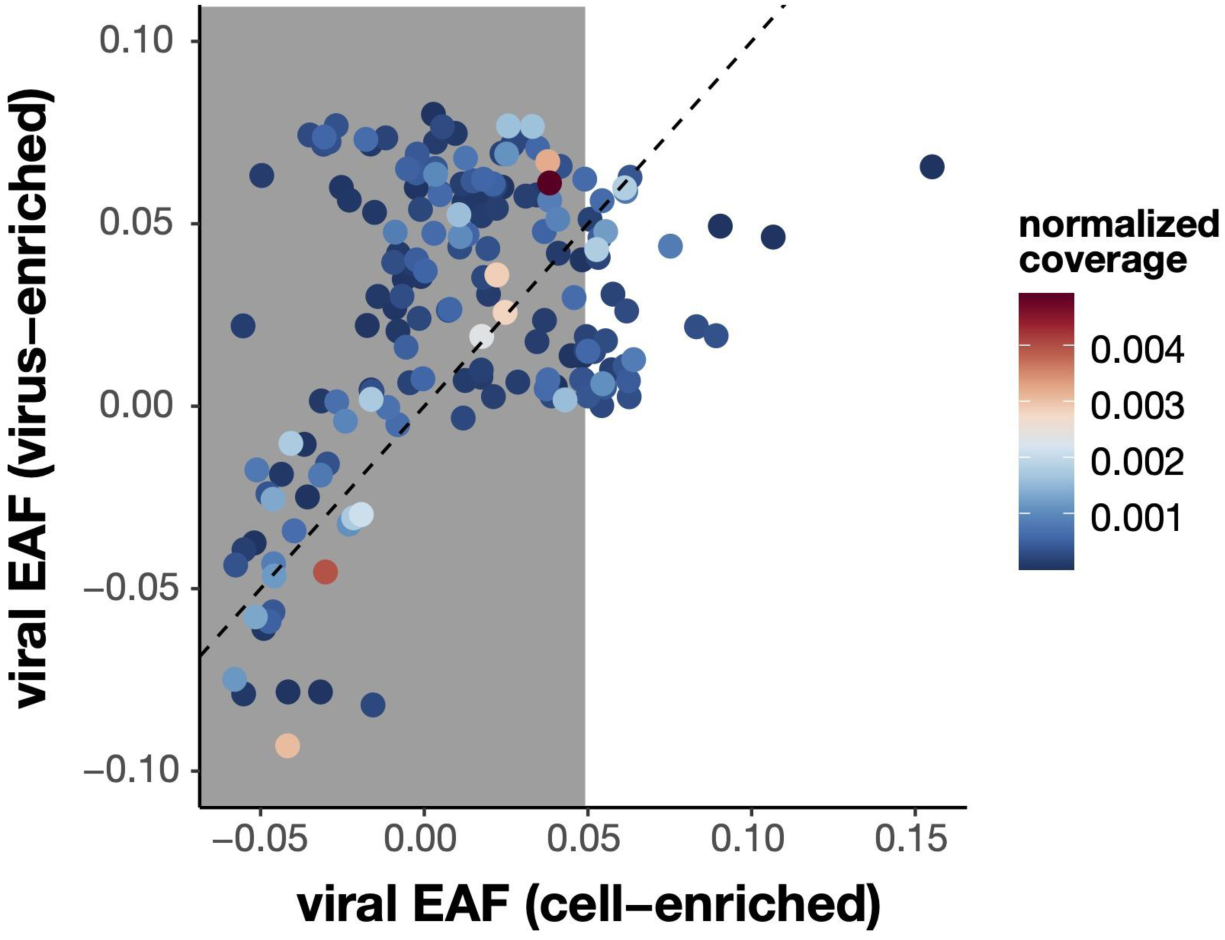
Carbon cycling activity as determined by the ^13^C Excess Atom Fraction (EAF) of high-confidence population genomes in the virus-enriched and cell-enriched size fractions. Viral population genomes are colored by their relative abundance (normalized interquartile coverages) in the virus-enriched sample. The shaded area represents insignificant carbon cycling activity. The dashed line represents the 1:1 ratio.

For each viral population genome, we calculated its approximate of turnover time, τ, during the incubation period using the ratio of its isotopic enrichment in the virus-enriched size fraction, *f*_*vir*_, relative to that in the cell-enriched size fraction, *f*_*cell*_, using the formula, *τ* = −*t*/(*ln*(*1* − *f*_*vir*_/*f*_*cell*_)). The most active viral population showed an EAF of 0.066 in the virus-enriched size fraction and 0.155 in the cell-enriched size fraction, giving a *f*_*vir*_/*f*_*cell*_ ratio of 0.426 and a turnover time of 12.6 days using the above equation. The second-most-active viral population was estimated to have a *f*_*vir*_/*f*_*cell*_ ratio of 0.43 giving a turnover time of 12.5 days. The third-most-active viral population had a *f*_*vir*_/*f*_*cell*_ ratio of 0.55 giving it a turnover time of 8.8 days. The above approximation for τ assumes the value of *f*_*cell*_ reaches a steady state very quickly, so we also developed a non-steady-state model to estimate τ from the rates of autotrophic production and viral pool size determined from model fits to the POC, PO^13^C and EAF data (see Supplemental). The non-steady-state model indicates that autotrophic production was 21.7% of the total prokaryotic production and predicts turnover times ranging from 4.9 to 7.2 days.

Based on these calculations, we estimate the turnover rates of viruses targeting chemoautotrophs in this environment at 5 to 13 days. This range of turnover rate estimations likely reflect lineage-specific variability in viral production and decay driven by biological characteristics and environmental conditions. This range of estimates is consistent with a general expectations of lower viral decay rates, which contribute to lower turnover rates, in the absence of light-induced degradation^46^, and with previous estimates of turnover rates ranging from 0.036 - 30 days in aquatic habitats^45^. Whereas previous calculations of viral turnover was restricted to bulk, community-wide estimates^45^, our population-specific estimates of viral turnover rates can be utilized to develop and improve ecosystems models in complex communities.

## Cross-feeding and multi-trophic interactions

As a typical consideration for tracer-based experiments, cross-feeding can allow for labeled nucleotides into other non-targeted trophic groups. In our case, cross-feeding (heterotrophic assimilation of ^13^C-organic carbon, decomposition of viruses^47^, and/or anaplerotic reactions) could have allowed heterotrophic cells and their viruses to incorporate ^13^C into their genomes. Common aquatic heterotrophs did not show significant label incorporation (Fig. 2), indicating undetectable cross-feeding in our incubation timeframe of 7 days. Our results show that SIP incubations can prevent cross-feeding while targeting microbial populations responsible for a specific process.

We observed no evidence of eukaryotic grazing of chemoautotrophs, as no environmental sequences were recovered from known eukaryotic protists. 0.00092% of reads from the environmental sample were identified as eukaryotic, all of which were classified as fungi. The majority (0.00075%) were classified *Malassezia restricta*, which has been observed across marine water column samples, deep-sea hydrothermal vents, sediments, and anoxic environments^48^. Given this lack of evidence for eukaryotic grazing, we postulate that viral lysis is the predominant mechanism that recycles chemoautotrophic biomass in this environment.

## Metabolic pathways and primary productivity

A metagenome-assembled-genome (MAG)-level analysis of 158 medium to high-quality MAGs (Fig. S8) indicated that at incubation temperatures of 20°C, the primary mechanisms of carbon fixation were the reductive tricarboxylic acid (rTCA) and Calvin-Benson-Bassham (CBB) pathways. While the WL pathway was recovered from sulfate-reducers (*Desulfosarcina*, *Desulfobacula*), it did not appear to contribute to carbon fixation (Fig. 2), consistent with previous reports of sulfate-reducing bacteria utilizing this pathway in reverse for anaerobic oxidation of organic matter^49^. The carbon cycling activity of MAGs with predicted rTCA pathways (*Sulfurimonas*) were similar to those with CBB (*Thiomicrorhabdus*, *Hydrogenovibrio*). Assuming similar rates of cellular turnover amongst these genera, these results suggest that carbon fixation efficiency amongst populations with rTCA was similar to those with CBB in this environment.

A gene-level analysis, normalized to a prokaryotic single-copy marker gene, indicated that sulfur oxidation genes were enriched in cellular populations with significant carbon cycling activity (Fig. S9). Taxonomic annotations of contigs with sulfur oxidation genes confirmed that these sequences were unique to genera with high carbon cycling activity (*Thiomicrohabdus*, *Hydrogenovibrio*, *Sulfurimonas*, and *Sulfurovum*). Hydrogen oxidation genes such as hydrogenases were not enriched in populations with significant carbon cycling activity (Fig. S9). Our results show that, although these active genera have been reported to oxidize both sulfur and hydrogen, sulfur oxidation is the predominant mechanism of chemoautotrophic primary production in this ecosystem. Taken together, these results demonstrate the effectiveness of qSIP-metagenomics in linking microbial diversity across multiple scales (trophic levels, populations, pathways, and genes) to key ecosystem functions and processes in complex environmental samples.

## Conclusions and future directions

Linking microbial diversity (via genomes) to key ecosystem processes (e.g., carbon cycling) remains a major challenge in environmental microbiology. In this study, we addressed this challenge by combining ^13^C-DIC SIP with metagenomics to track the flow of carbon from the environment into cellular and viral genomes. We conducted these experiments targeting the aphotic chemocline of a marine-derived meromictic coastal pond as a model system to identify key processes governing carbon cycling in chemoautotrophic communities. We found that primary production in this ecosystem is driven by highly active low abundance genera that could have likely been overlooked using metagenomic approaches alone (*Thiomicrorhabdus*, *Hydrogenovibrio*, *Sulfurimonas*, *Sulfurovum*). Despite the absence of informative taxonomic or functional annotations, we linked novel viruses to these key chemoautotrophs using similarities in the isotopic signatures of viral and host genomes. Through this reference-independent approach, we found evidence for the virus-mediated recycling of chemoautotrophic biomass through the production of viral particles. Given the broad distributions of these key sulfur-oxidizing chemoautotrophs across marine and terrestrial environments^2,4,5,14–18^, our results highlight a previously overlooked component microbial carbon cycling that we postulate has global-scale impacts.

We demonstrated the ability to estimate population-specific rates of viral turnover using isotopic enrichment data and anticipate that this data will be foundational for building and validating novel ecosystems models. This data will also be useful in benchmarking *in silico* metagenomic programs to enable more accurate and high-throughput predictions of virus-host linkages, as well as cross-validating with wet lab approaches, such as single-cell and hi-C sequencing, to link viruses to their hosts. Furthermore, we will leverage this reference-independent approach to develop new methods to predict the biogeochemical impacts of microbial populations and discover novel pathways underpinning key ecosystem processes. We reason that our approach is particularly useful in understudied environments to provide critical links between microbial diversity and key ecosystem processes across multiple biological scales (trophic levels, populations, pathways, and genes).

## Supporting information

Luo et al. 2025 Supplemental Information

Luo et al. 2025 Model Equations

Luo et al. 2025 Supplemental Tables

## End notes

## Acknowledgements

We would like to acknowledge Alex Worden, Gretta Serres, Sabrina Elkassas, Cynthia Becker, and Amy Apprill for discussions and/or methodology relevant to this manuscript. The metagenomic data generation, data analyses, and manuscript was supported by the University of North Carolina at Charlotte (startup funds to EL). The wet lab work was supported by the Woods Hole Oceanographic Institution (Weston Howland Jr. Postdoctoral Fellowship to EL), the National Oceanic and Atmospheric Administration (NA19OAR4320072 subaward 0007525/102212019 to JAH), and the US National Science Foundation (OCE-1947776 to JAH). JJV was supported by the Simons Foundation (549941FY22). BEB was supported by the US National Science Foundation (PRFB2010963). GT was supported by the US Department of Energy (BER GSP “Microbes Persist” SCW1632 and contract DE-AC52-07NA27344).

## Author contributions

EL conceptualized, supervised, and acquired funding for the study with input from JAH. EL and BEB collected the samples with help from JJV and JAH. EL conducted the experiments and wet lab analyses. EL, NDP, TJR, and JJV conducted the data analyses. EL wrote the manuscript with input from co-authors. The authors declare no competing interests.

Supplementary Information is available for this paper. The datasets generated for this current study will be made available on FigShare with publication. Correspondence and requests for materials should be addressed to

## Methods

### Sampling site

Water samples for microbial analysis were collected at 10m from Siders Pond, a year-round marine-derived coastal meromictic kettle hole in Falmouth, Massachusetts (41.549006, - 70.622039°), typical of historically glaciated coastal regions. Its stratification is due to inputs of both freshwater and seawater, and the lack of seasonal mixing leads to a shallow chemocline that separates the photic, oxygenated upper freshwater from the aphotic, anoxic bottom saline water (Fig. S1). Anoxic conditions (<0.2mg/L) were observed below 8.5m (Fig. S2). An increase in hydrogen sulfide in the bottom saline water was observed at 8-10m^50,51^. We chose a sampling depth of 10m to target chemoautotrophic sulfur oxidizers at the chemocline.

### SIP incubation

Water from 10m deep in Siders Pond was collected on Nov 11th, 2021 using a handheld pump attached to a YSI probe measuring photosynthetically active radiation, dissolved oxygen, and salinity (Fig. S1). 1L of this environmental sample was filtered through 0.22μm filters (Millipore-Sigma Sterivex SVGP01015) and then through 0.02μm filter (Whatman Anotop WHA68092102) to respectively collect the cell-enriched and virus-enriched size fraction as environmental controls. Six samples were incubated at 20°C in the dark in 1L glass sealed bottles that were pre-purged by removing air, filling with nitrogen gas, and purged again prior to filling with sample fluid. Each 1L bottle was dosed with 7.33mL of 400mM ^12^C-sodium bicarbonate (^12^C controls) or ^13^C-sodium bicarbonate (^13^C labeled treatments) for a final concentration of 2.9mM. The concentration of DIC at 10 m in Siders Pond was measured at 4.11 mM on 6-Oct-2021 by students in MBL’s Semester in Environmental Science program, which would produce a 41.3% ^13^C isotopic enrichment of DIC in the ^13^C treatment. The incubations were terminated by sequential filtration through 0.22μm and 0.02μm filters to respectively collect cell-enriched and virus-enriched samples at the following timepoints: 1 day (^12^C + ^13^C), 3 days (^13^C), 5 days (^13^C), and 7 days (^12^C + ^13^C).

### Label incorporation during incubation timeframe

50mL of sample was filtered onto combusted 0.45μm glass fiber filters, washed with 50mL of 1x phosphorus-buffered saline, dried in a 50°C oven overnight in individual sterile petri plates, and analyzed on gas chromatography mass spectrometry for quantification of isotopic enrichment. Filters were analyzed at the Marine Biological Laboratory Stable Isotope Laboratory for δ13C using a Europa 20-20 continuous-flow isotope ratio mass spectrometer interfaced with a Europa ANCA-SL elemental analyzer (Sercon Ltd., Cheshire, UK). From these results, the day 7 incubation (^12^C/^13^C pair) was chosen for downstream molecular analyses.

### DNA extraction and density-gradient centrifugation

DNA was extracted from the virus-enriched (0.02μm filters) and cell-enriched (0.22μm filters) samples from the environmental control and 7-day ^12^C/^13^C incubation pair using the Masterpure extraction kit (Lucigen MC85200), yielding 1.8 - 8.4μg and 20 - 25μg of DNA respectively from the virus-enriched and cell-enriched samples. A major challenge of qSIP-metagenomics is generating sufficient DNA biomass to enable successful density-fractionation and sequencing of multiple density fractions per sample. This consideration is particularly challenging for low-biomass samples, such as the viral-particle size fraction that we targeted in this study. Here, we show that density-fractionation of virus-enriched samples (0.02-0.22μm) and quantification of viral activity can be successfully achieved with 1L of aquatic samples.

Both virus-enriched and cell-enriched samples from the ^12^C and ^13^C treatments were density-fractionated as previously described^52^ with the following modifications. 500ng of input DNA was utilized for the virus-enriched samples, and 1000ng of input DNA was utilized for the cell-enriched samples. The input DNA was diluted using gradient buffer to 700μL total, added to 4.8mL of 1.77g/mL cesium chloride solution, and individually loaded into 5.1mL centrifuge tubes (Beckman Coulter Quick-Seal). Virus-enriched DNA was ultracentrifuged at 44,000g for 5 days (Beckman Coulter Optima XE), and cell-enriched DNA was separately ultracentrifuged at 44,000g for 3 days, both in a VTi 65.2 vertical rotor. Each sample was separated into 12 density fractions of ∼400uL each. The density of each fraction was measured using a hand-held refractometer calibrated with MilliQ water to nD-tC at 20°C = 1.3330, and converted using the equation (nD-tC at 20°C)*10.9276 −13.593 as previously described^53^. DNA from each fraction was precipitated using PEG solution^52^ and 75mg of Glycoblue (ThermoFisher AM9516). DNA recovered from each density fraction was quantified using Picogreen (ThermoFisher P11496) and qPCR (Fig. S3). 792 – 15217 ng of DNA was recovered in the cell-enriched and virus-enriched samples, enabling successful density fractionation of metagenomic DNA across treatments and size fractions. All samples showed consistent, distinct peaks in DNA density between ^12^C and ^13^C treatments that indicated ^12^C-DIC and ^13^C-DIC label incorporation (Fig. S3). 3 -6 of the heaviest and lightest fractions, which contain the lowest amount of DNA, were pooled to a minimum of 25ng DNA, resulting in 5 density fractions per sample for sequencing (Fig. S3) as previously described. DNA libraries were prepared using the Illumina KAPA HyperPrep kit and sequenced on the Illumina Novaseq X plus 10B platform.

### Read processing and metagenomic assembly

Raw reads were trimmed using two passes through BBDuk^54^ v39.01 to remove Illumina adapters and phiX with the following parameters: ktrim=r k=21 mink=11 hdist=2 tbo tpe for adapter removal during the first pass and k=27 hdist=1 qtrim=rl trimq=17 cardinality=t mingc=0.05 maxgc=0.95 for phiX removal during the second pass. Reads from cell-enriched (0.22μm) samples were pooled and separately assembled in three groups (environmental control, ^12^C, and ^13^C) using MEGAHIT^55^. Virsorter2 v.2.2.4^56^ was used to identify putative viral contigs in cell-enriched assemblies. QC’ed reads were then mapped using Bowtie2^58^ 2.5.1^57^ to these putative viral contigs in their corresponding assemblies to identify viral reads within cell-enriched samples. Viral reads from the three cell-enriched assemblies were pooled with the corresponding reads from the three virus-enriched (0.02μm) samples (environmental control, ^12^C, and ^13^C) and co-assembled in metaSPAdes with a minimum contig length of 1.5 kb. QUAST^58^ v5.2.0 was used to assess assembly quality.

### Viral population genomes

Putative viral contigs were identified from the viral assemblies using Virsorter2 v2.2.4^56^ retaining 48,925 putative viral contigs of >5kb from all categories across the three assemblies. Putative viral contigs were then dereplicated by clustering at >95% ANI across >50% of the shorter contig using anicalc.py and aniclust.py scripts^59^, resulting in 31,488 putative viral population genomes. CheckV ^59^ v1.0.1 was used to assess completeness. High-confidence viral population genomes were identified as previously described^60^, and remaining contigs that did not pass through the workflow were retained only if they contained one or more viral structural genes. Temperate phages were identified as previously described^61^. The VC ratio for each viral population, the ratio for its relative abundance in the virus-enriched fraction relative to that in the cell-enriched fraction, was calculated as previously described^19^.

### Viral taxonomic and functional annotation

Viral taxonomic assignments were performed using two complementary approaches: (i) marker-based classification via geNomad^62^; (ii) protein similarity searches against reference viral databases, including NCBI NR and IMG/V4. For the marker-based assignment, the geNomad^62^ v1.7.0 pipeline (genomad end-to-end) was used to classify sequences using taxonomically informative protein profiles. To assign taxonomy based on similarity, open reading frames were predicted using Prodigal^63^ v.2.6.3 with the parameter “-p meta”. The predicted ORFs were then compared to viral proteins in NCBI NR^64^ (retrieved in 2025-02-14) and IMG/V4^65^ using used DIAMOND^66^ v.2.9. Each viral polation genome was assigned to the lowest common taxonomic rank supported by at least 50% of its annotated proteins at >60% AAI. Metacerubus^67^ was used for functional annotation, which includes the KEGG^65^, COG^69^, VOG^70^, PHROG^71^, and PFAM^72^ databases. Putative prophages were identified using marker genes as previously described^19,73^.

### Virus-host linkage

Viral contigs were linked to their hosts through CRISPR spacer mapping using the CRISPRCASFinder^74^ and CRASS^75^ programs based on 100% alignment across 100% of the spacer, as well as mapping to prophages at >95% ANI across >1kb to a cellular contig that is at least twice the length of the viral contig. The taxonomy of the linked hosts were identified through Kaiju v1.10.1^76^.

### Metagenomic-assembled genomes (MAGs)

Assemblies from the cell-enriched (>0.22μm) samples were run through the Binning and Bin_refinement modules of MetaWRAP^77^ to construct MAGs. Within the Binning module, MaxBin2^78^, MetaBAT2^79^ and CONCOCT^80^ were used for curating the initial bin sets. The Bin_refinement module was used to refine the bin sets into a consensus bin set for each assembly for a total of 227 bins at ≥50% complete and <10% contamination. MAGs were then dereplicated using dRep^81^ into one consensus bin set of 158 MAGs (Fig. S7). CheckM^82^ v1.1.3 was used for both the Bin_refinement module and dRep to evaluate MAG completeness and contamination based on prokaryotic lineage-specific marker genes.

### Cellular functional and taxonomic annotation

Kaiju^76^ (default setting) was used for taxonomic identification of cellular contigs. The genome Taxonomy Database Toolkit^83^ v. 2.1.1 (GTDB) was used for taxonomic identification of MAGs. DRAM^84^ and Metacerubus^67^ were used for the functional annotation of MAGs, which includes the KEGG^65^ and COG^69^ databases. The keggLink function from the KEGGREST^85^ package on R was used to pull all KO numbers for each carbon fixation pathway from the KEGG database. MAGs were predicted to contain a carbon fixation pathway if they contained all key enzymes as well as ≥50% of genes in the complete KEGG pathway, consistent with a previous report^6^. Normalized gene counts, approximating copies per genome, was calculated by dividing the number of genes in that functional category by the number of universal single-marker genes (COG0012/KO6942).

### Quantifying taxon-specific isotope incorporation

Bowtie2^57^ and samtools^86^ were used in conjunction with Anvi’o^87^ v.7.1 to calculate interquartile coverage for each contig, which reduces the impact of conserved or hypervariable regions in respectively over/underestimating coverage calculations. R^88^ ‘decostand’ function of the vegan^89^ package v2.6-10 was used to convert interquartile coverages for each contig into relative abundances. The excess atom fraction (EAF) of 13C within each contig was calculated as previously described^90^ with the following modifications. The total genomic copy per μL of contig i within density fraction k was calculated by multiplying the total number of genome copies (f; determined by either qPCR ratio for the cell-enriched fractions or DNA yield ratio for the virus-enriched fractions) of density fraction (k) by the relative abundance (R) of contig i within density fraction k. Additionally, instead of calculating the GC content using the regression formula, we calculated GC content of each contig by using the program seqkit^91^ with the following parameters: ‘seqkit fx2tab –name –only-id –gc’. To improve accuracy in our calculations, we retained only contigs that were present in three or more density fractions in both treatments. To identify populations that demonstrated significant carbon incorporation, we plotted the calculated EAF values against the ranked EAF values (numerically ranked from lowest to highest position). A segmented linear regression was then run against the resulting spline function to identify a break point at which ^13^C incorporation can be inferred^27^. The function segmented from the R segmented package^92^ was used to identify the breakpoint, which was identified at rank 9067.20 with a standard error of 3.28. Three times the standard error was added to this breakpoint to identify the EAF threshold of 0.049. Microbial populations with EAF calculations above this threshold were defined as having significant carbon cycling activity.

### Microbial richness in environmental sample and post-incubation

Cellular richness was estimated by calculating the number of non-redundant cellular contigs with non-zero interquartile coverage in the cell-enriched size fraction in the environmental sample, ^12^C, and ^13^C treatments. Viral richness was estimated by calculating the number of viral population genomes with non-zero interquartile coverage in the virus-enriched size fraction in the environmental sample, ^12^C, and ^13^C treatments.

### Identification of eukaryotic sequences

Eukdetect^93^ v1.3 “run all” and RiboTagger^94^ v0.8.0 were used to identify eukaryotic reads. Eukdetect identifies eukaryotic sequences based on a database of 521,824 microbial eukaryote marker genes, while RiboTagger identifies eukaryotic DNA by aligning reads to 18S rRNA database (v4, v5, v6, and v7 regions). For quantitative comparison among different metagenomics datasets, the relative abundance of eukaryotic species was calculated by normalizing the number of reads mapped to the identified species to million sequencing reads (reads per million) in individual metagenomes.

## Extended Data

**Figure S1.**
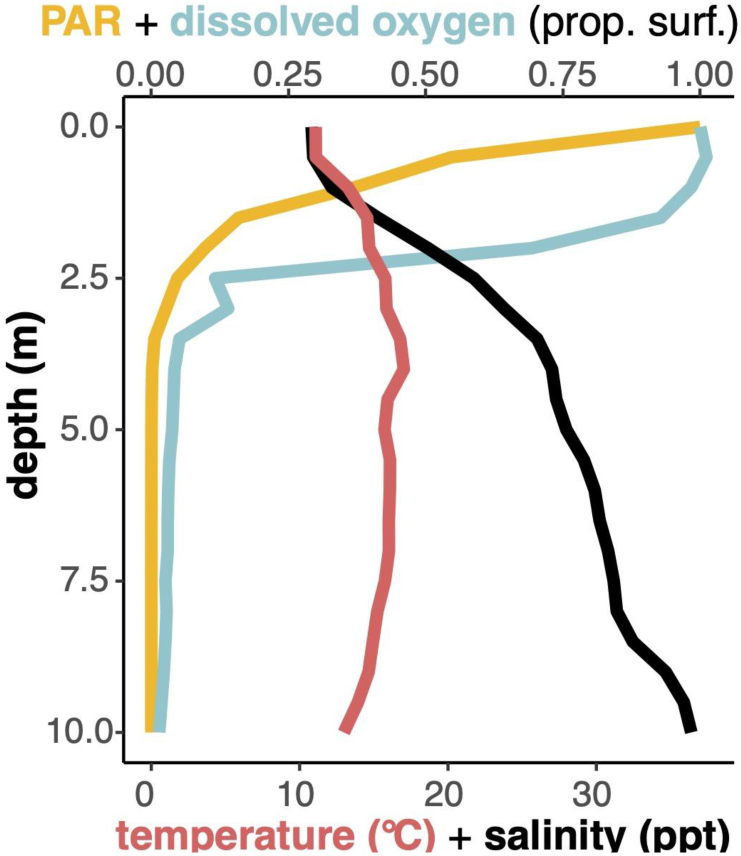
Depth profile of environmental metadata at Sider’s Pond associated with sample collection at 10m on November 11th, 2021. Photosynthetically active radiation (PAR) and dissolved oxygen levels are plotted as a proportion relative to the surface value (384μmol/m²/s and 9.95mg/L, respectively).

**Fig S2.**
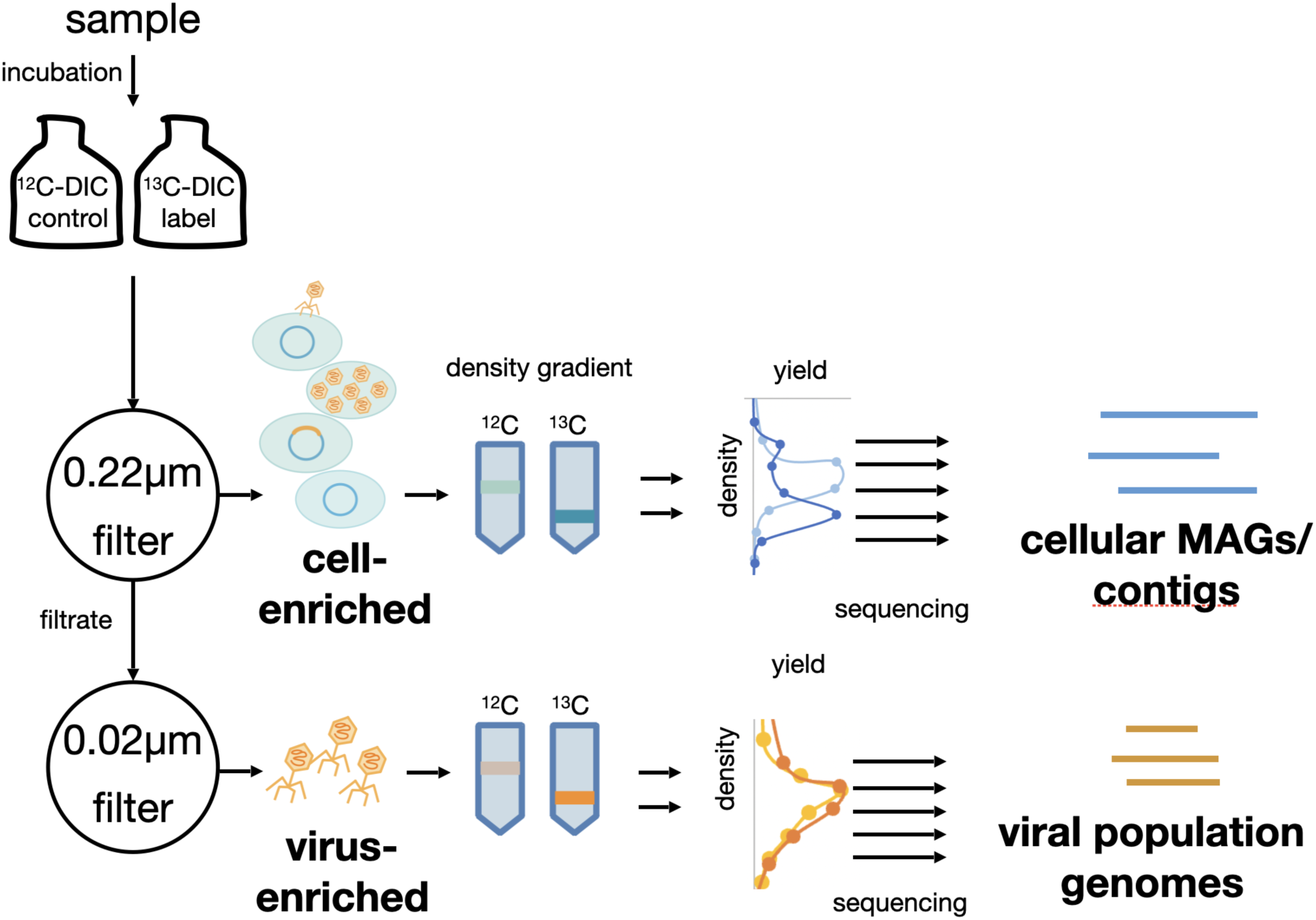
Incubation and qSIP-metagenomics workflow conducted in this study. DNA density and yield curves represent real data shown also in Fig. S3.

**Figure S3.**
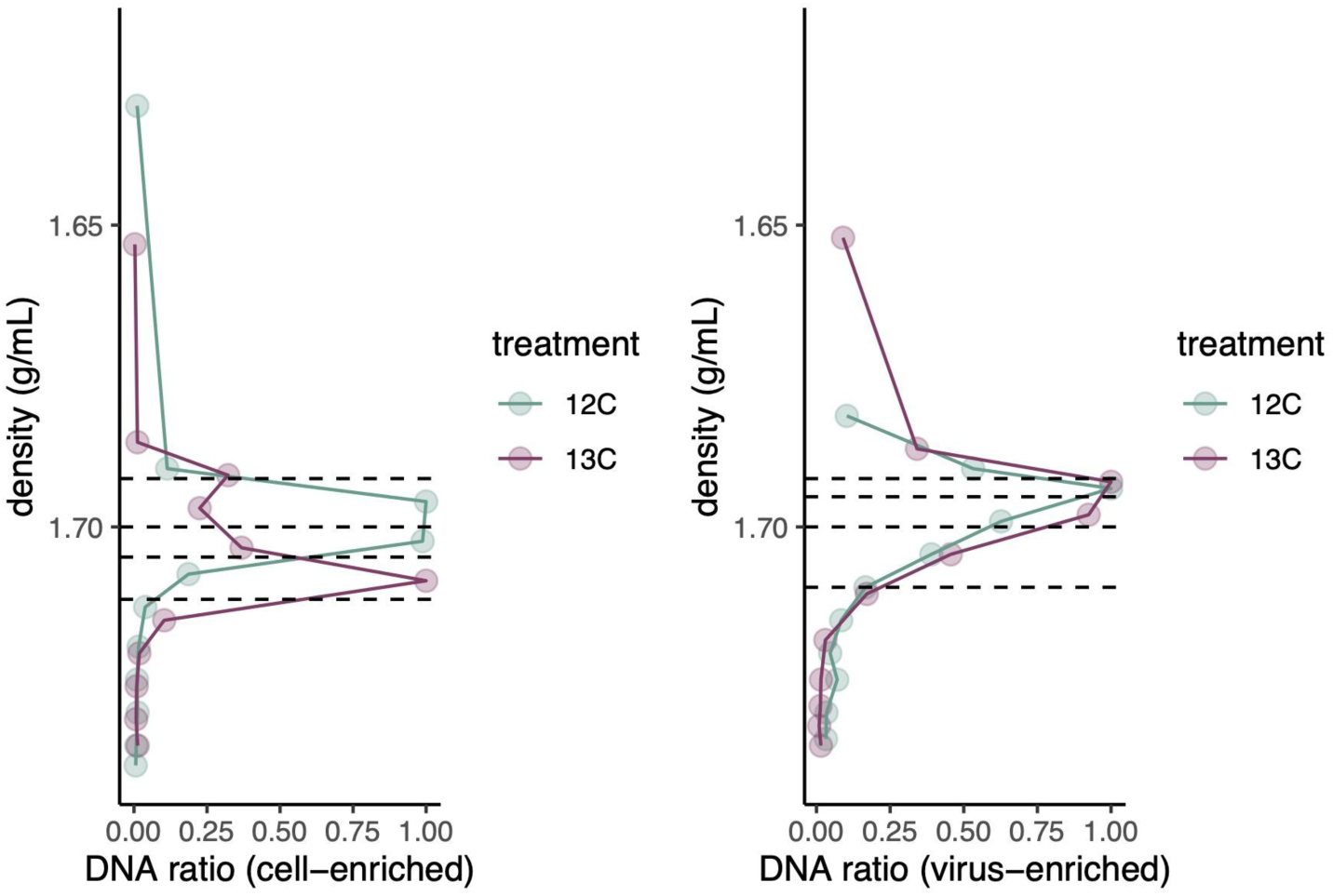
Density curves of cell-enriched (>0.22μm) and virus-enriched (0.02 - 0.22μm) samples, 7 days post-incubation. The dashed line represents separations between the five density fractions sequenced. 3-5 of the heaviest and lightest fractions were pooled prior to sequencing. The lightest fractions are omitted from visualization due to the standard inclusion of water in that fraction. The DNA ratio represents the qPCR yield in the cell-enriched size fraction (16S rRNA gene copy number, approximating prokaryotic DNA yield) and the total DNA yield in the virus-enriched size fraction (approximating total DNA yield) across the density gradient, normalized to the highest value across density fractions in the respective samples.

**Figure S4.**
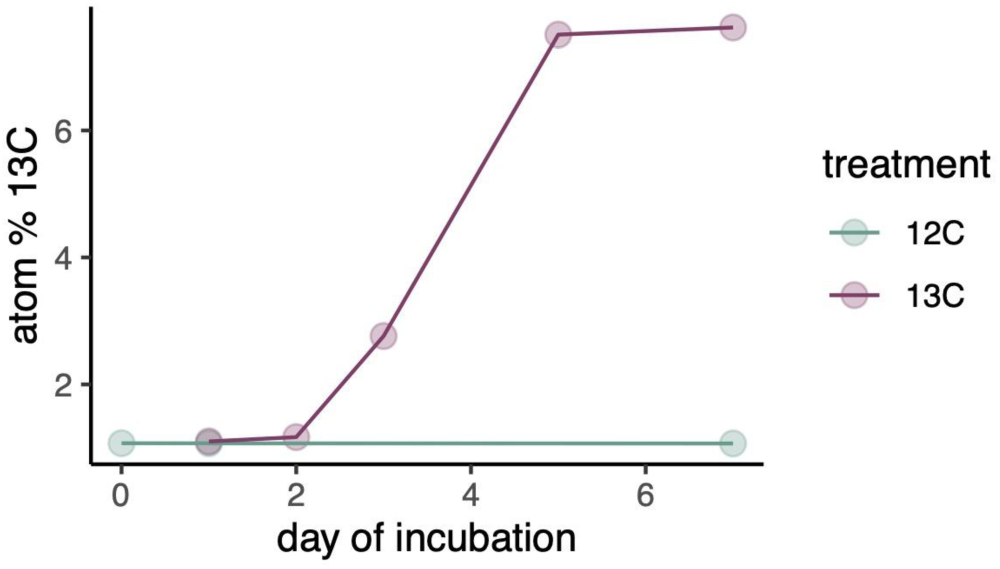
13C-isotopic enrichment in the >0.45μm particulate organic carbon fraction during the 7-day incubation period, as quantified by gas chromatography mass spectrometry.

**Figure S5.**
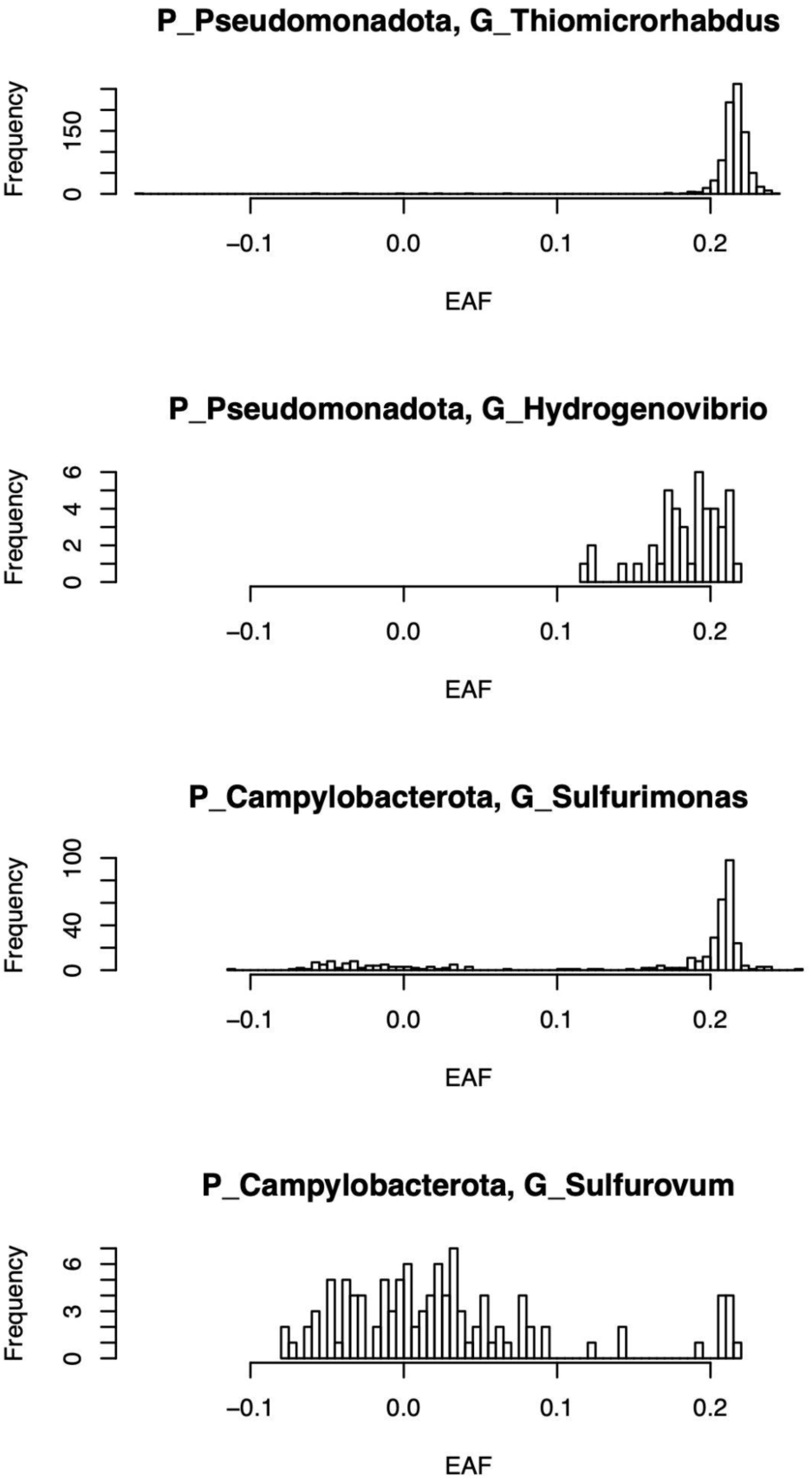
Histograms of Excess Atom Fraction (EAF) values for all contigs belonging to the four most active genera (P = phylum, G = genus), annotated with Kaiju for unbinned contigs and GTDB for binned contigs.

**Figure S6.**
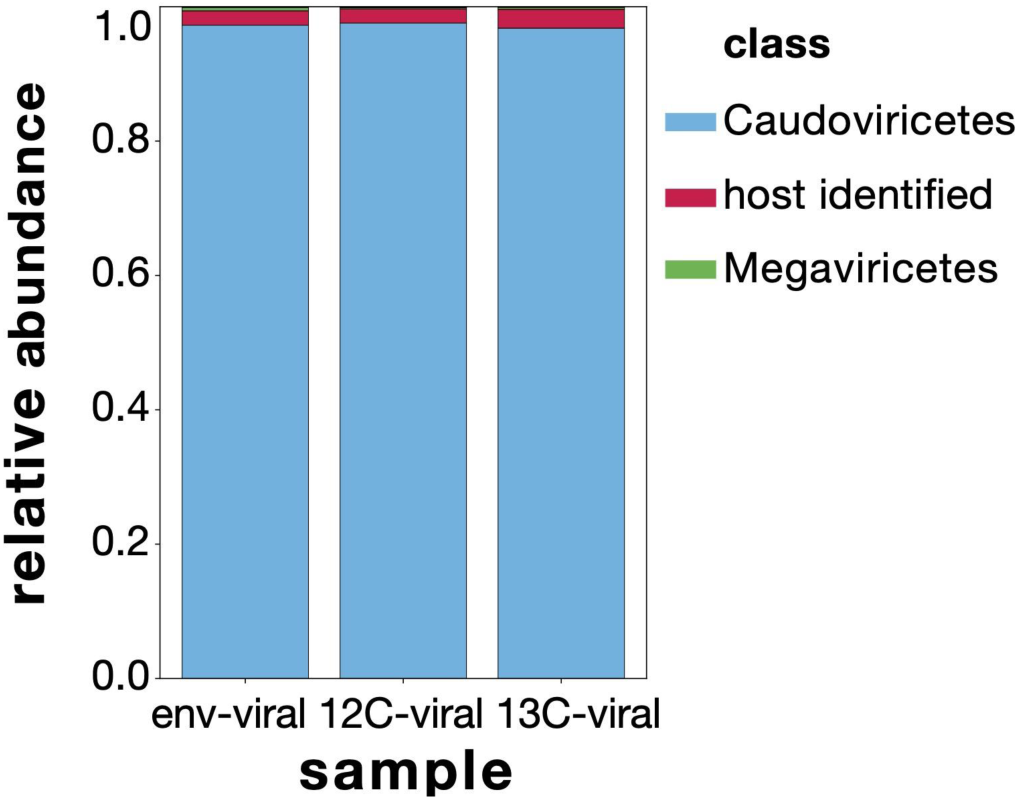
Relative abundances of viral assemblages (0.02 - 0.22μm) recovered from the environmental control (env-viral) and post-incubation (12C-viral, 13C-viral) samples. The relative abundances of viral populations are colored by taxonomic annotations from geNomad and reference databases at the class level (the most detailed taxonomic resolution that yielded annotations using these methods). Low-abundance annotations that cannot be visualized are not listed. Host-identified contigs through prophage and CRISPR spacer mapping are colored in red.

**Figure S7.**
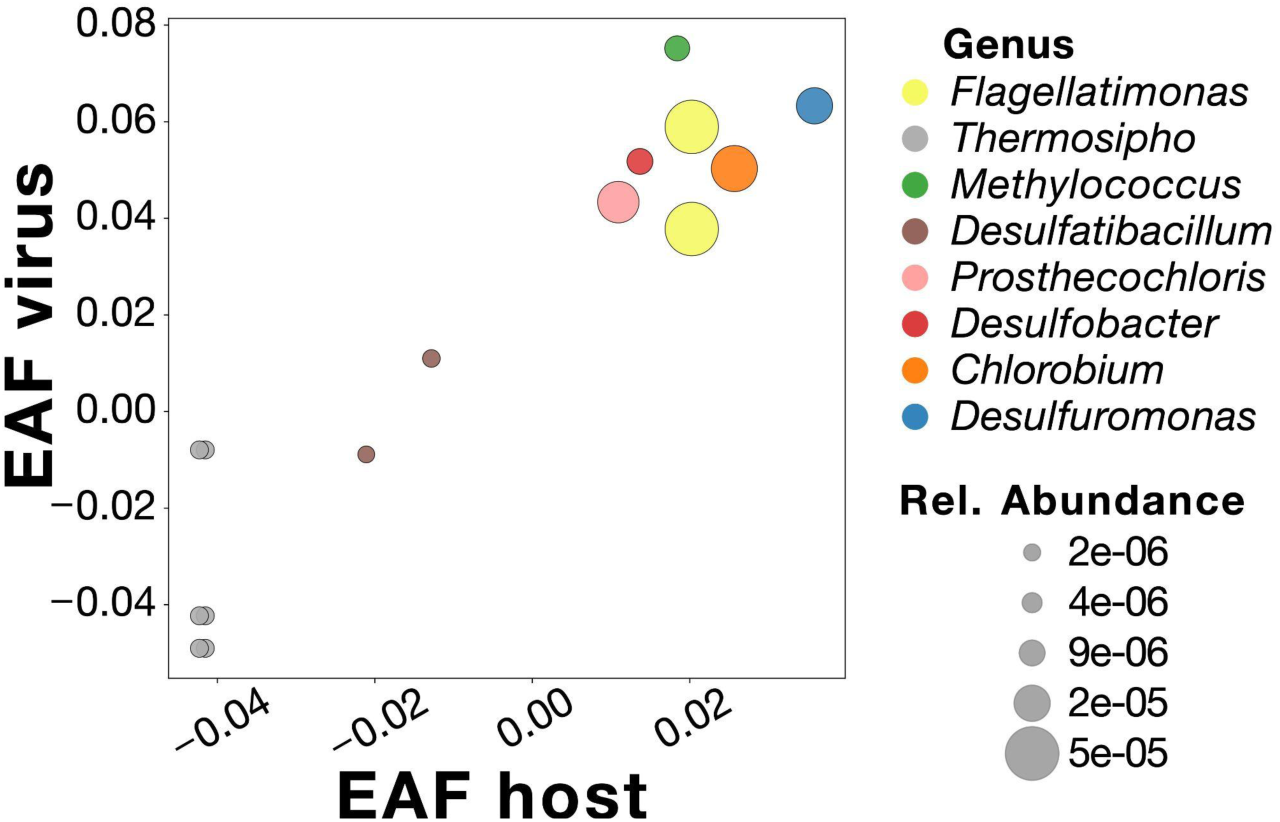
Carbon cycling activity as determined by 13C Excess Atom Fraction (EAF) of 15 high-confidence viral population genomes (y-axis) and their predicted cellular hosts (x-axis). Viral population genomes were linked to cellular contigs using in silico prediction based on prophage and CRISPR spacer mapping. Virus-host pairs are color-coded by the genus-level taxonomic assignment of the predicted cellular host contigs. The size of each circle corresponds to the host’s relative abundance in the environmental sample, as determined by the contig’s normalized interquartile coverage in the cell-enriched size fraction.

**Figure S8.**
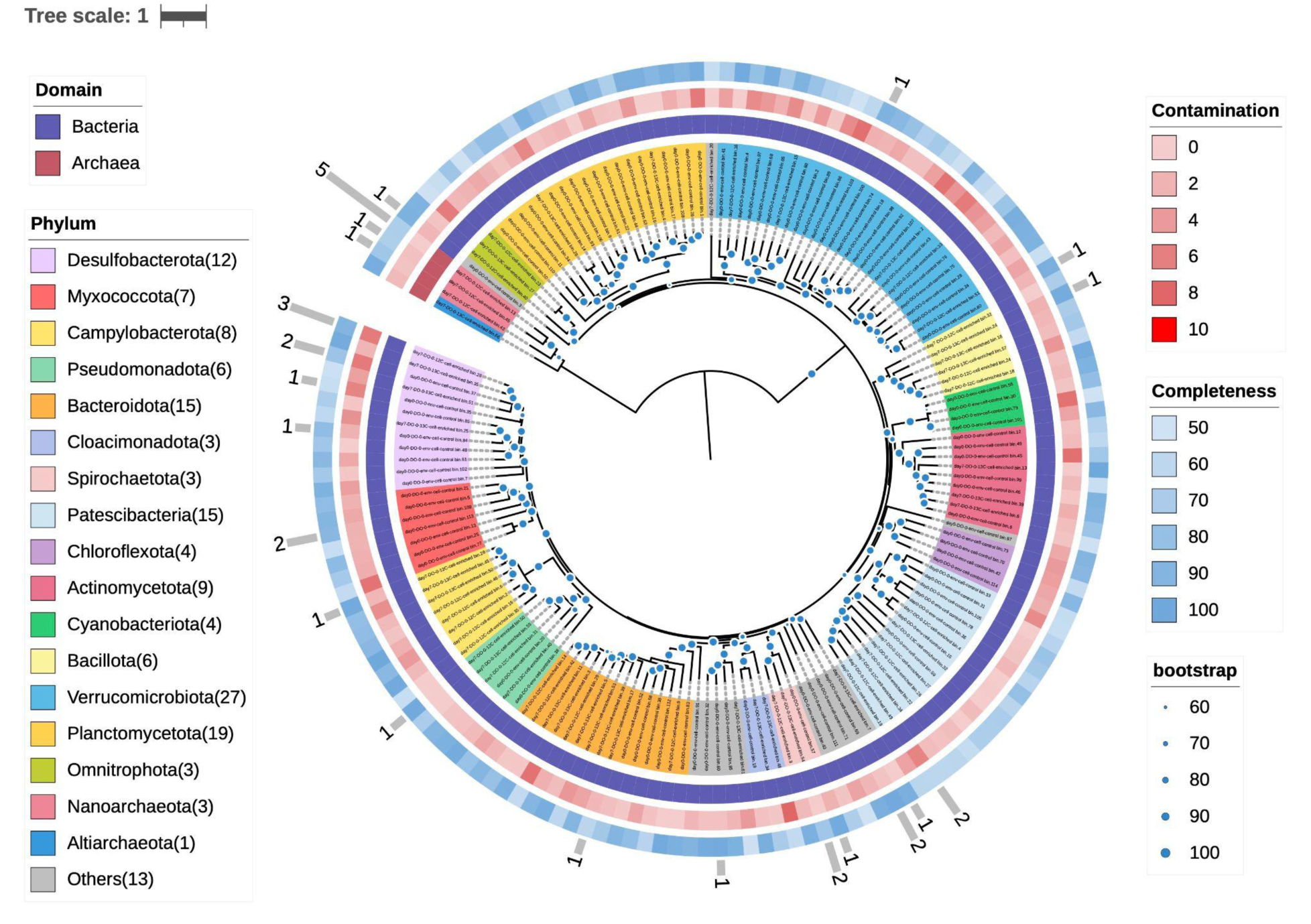
Phylogenetic tree of 158 non-redundant MAGs with >50% completion and <10% contamination, generated from IQtree, colored by phylum-level assignments as determined by GTDB-TK. The outer ring bar graph with numbers indicates the number of viral population genomes that were linked to each MAG through prophage and CRISPR spacer mapping.

**Figure S9.**
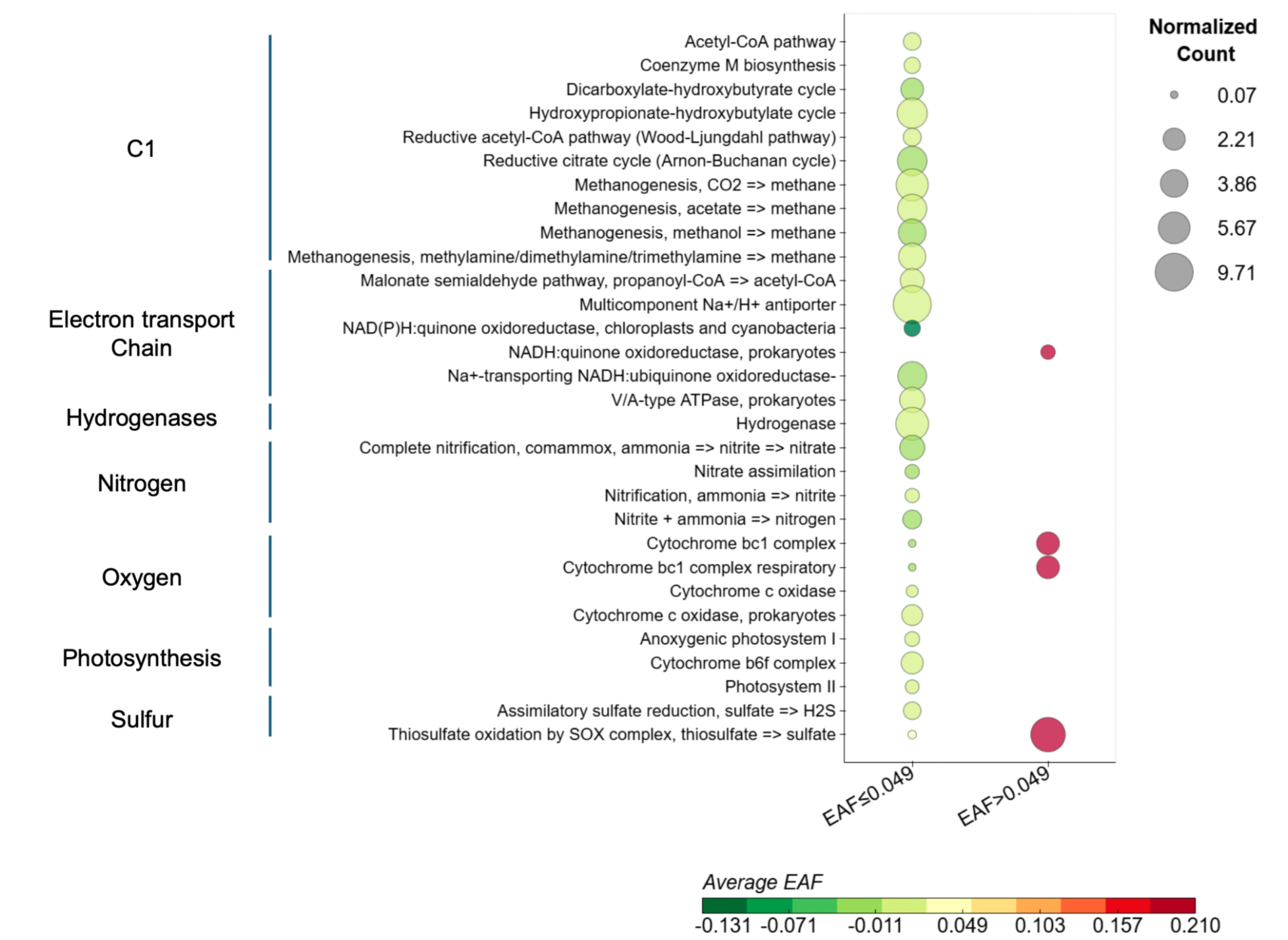
Differences in the predicted functional capacity of cellular sequences based on their carbon cycling activity. Cellular contigs are grouped on the x-axis and colored by whether they showed insignificant (EAF<=0.049) or significant carbon incorporation (EAF>0.049). The average EAF for each functional category was shown and color-coded on the scale bar below. The size of the circles represents the normalized gene count, calculated by number of genes in that functional category divided by the number of universal single-marker genes (COG0012/KO6942) found on the contigs in that group, as a proxy for gene copies per genome of that particular function in that group. Functional categories with a >=5-fold difference in the normalized gene count amongst the two cellular contigs groups (EAF>0.049 and EAF<=0.049) are shown.

