## Supplementary material for "Virus-mediated recycling of chemoautotrophic biomass": Luo et al. 2025 Supplemental Information

**Luo_et_al_SI_tables.xlsx**

Table S1. EAF values of cellular contigs. Columns are: contig, MAG (if binned), GC content, average density in the 12C treatment, average density in the 13C treatment, Excess Atom Fraction, and taxonomic assignment.

Table S2. EAF values of viral population genomes. Columns are: contig, GC content, nucleotides mapped from the virus-enriched environmental sample, nucleotides mapped from the cell-enriched environmental sample, VC ratio, log VC ratio, average density in the 12C treatment (cell-enriched sample), average density in the 13C treatment (cell-enriched sample), Excess Atom Fraction (cell-enriched sample), and taxonomic assignments.

Table S3. Species richness of cells and viruses in the environmental sample and post-incubation.

Table S4. EAF values of viral population genomes. Columns are: contig, GC content, normalized interquartile coverage in the virus-enriched environmental sample, normalized interquartile coverage in the cell-enriched environmental sample, nucleotides mapped from the virus-enriched environmental sample, nucleotides mapped from the cell-enriched environmental sample, average density in the 12C treatment (cell-enriched sample), average density in the 13C treatment (cell-enriched sample), Excess Atom Fraction (cell-enriched sample), average density in the 12C treatment (virus-enriched sample), average density in the 13C treatment (virus-enriched sample), Excess Atom Fraction (virus-enriched sample), and taxonomic assignments.

**Luo_et_al_ModelEquations.docx**

Microsoft Word file that describes the model equations used to calculate viral pool turnover rates.
