## Supplementary material for "Virus-mediated recycling of chemoautotrophic biomass": Luo et al. 2025 Model Equations

### Simple POC with ^13^C enrichment model

Mass balance around POC pool is

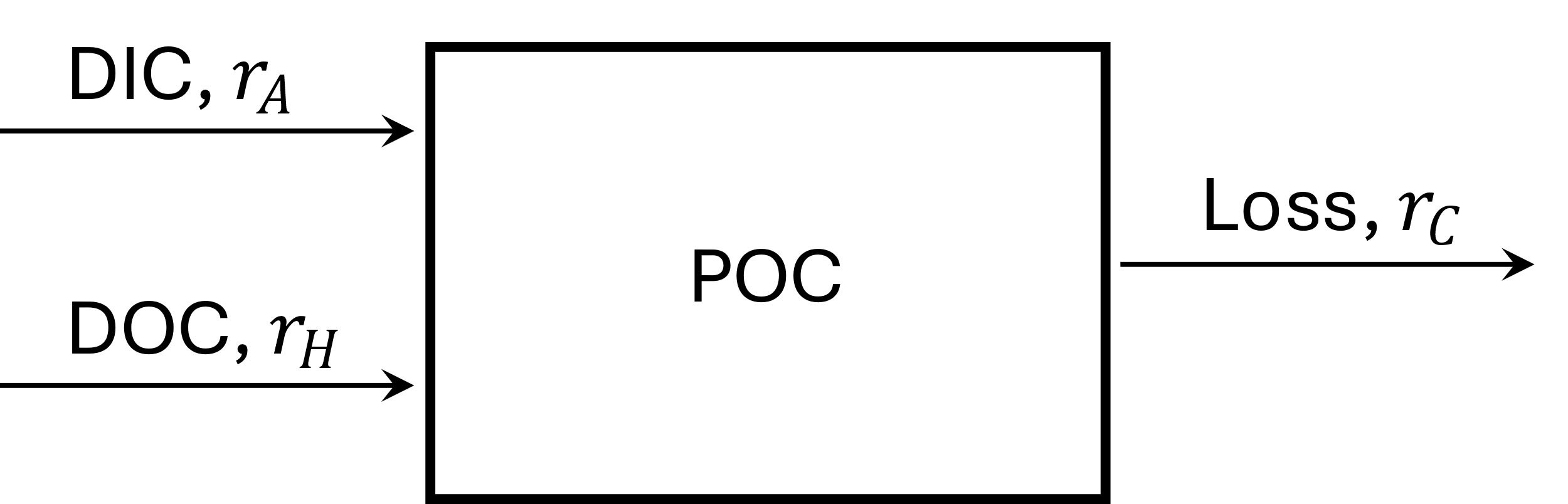

Where $r_{A}$ is the rate (μM d^-1^) of DIC fixation into POC by chemolithoautotrophs and $r_{H}$ is the rate of POC production from heterotrophs from DOC. POC can be lost back into DIC (via respiration) or to DOC at a rate of $r_{C}$ (but this does not change the ^13^C of the DOC pool). A mass balance around PO^13^C can be constructed, where $f_{A}$ is the fraction of DIC that is ^13^C, which is thought to be 0.413 and $f_{H}$ is the fraction of DOC that is ^13^C, which is believed to be just the natural abundance level (0.011). The governing equations for the concentrations of POC (µM) and PO^13^C (µM) are given by,

$$\frac{dC_{POC}\left( t \right)}{dt}=r_{A}+r_{H}-r_{C}; C_{POC}\left( 0 \right)=145 \mu M$$

(1)

$$\frac{d{{}^{13}C}_{POC}\left( t \right)}{dt}=f_{A}r_{A}+f_{H}r_{H}-\frac{{{}^{13}C}_{POC}\left( t \right)}{C_{POC}\left( t \right)}r_{C}; {{}^{13}C}_{POC}\left( 0 \right)=f_{H}C_{POC}\left( 0 \right) \mu M$$

(2)

The fraction of ^13^C in the POC, $f_{C}(t)$, is $\frac{{{}^{13}C}_{POC}\left( t \right)}{C_{POC}\left( t \right)}$. A simple constant model for the, $r_{A}$, $r_{H}$, and $r_{C}$ did not fit the $C_{POC}\left( t \right)$ and $f_{C}(t)$ observations very well, and exponential models for the rates (such as $Ae^{\mu_{A}t}$) fit the data well, but produced rates that were too high by day 7 (~1500 µM d^-1^). Simple linear rate models given by,

$$r_{A}=At+A_{0}; r_{H}=Ht+H_{0}; r_{C}=Ct+C_{0} ,$$

(3)

were a good compromise and able to fit the data well and produced rates that were reasonable, where the units for $A$, etc are µM d^-2^ and for $A_{0}$, etc. are µM d^-1^. The parameters $A, A_{0}, H,$ etc, were determined by fitting the modeled POC and $100\times f_{C}(t)$ to the observations via least squares subject to the ODE constraints, Eqs (1) and (2). It was assumed that the standard deviation of the measures was ±5 µM for POC and ±0.1% for PO^13^C%. Model fit (blue lines) to observations (red circles) are shown here,

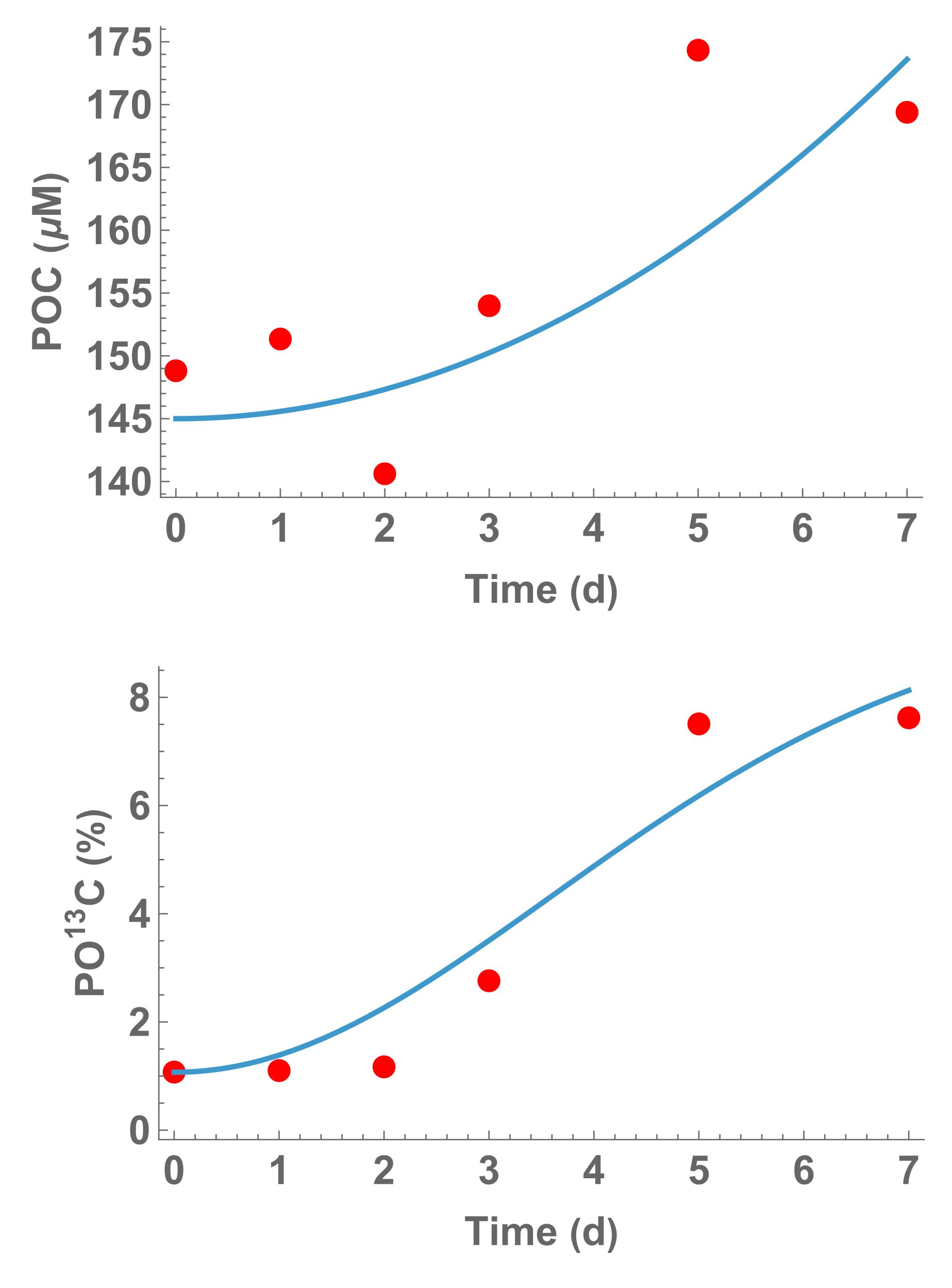

For a DI^13^C labeling of 41.3%, the parameters values found were $A=2.299,A_{0}=0,H=8.272,H_{0}=0,C=9.386 \mu M d^{-2} ,\text{ }\text{and}\text{ }C_{0}=0.052 \mu M d^{-1}$. The fraction of the POC production that came from chemoautotrophs was 21.7% based on these rates. The rates of $r_{A}$ and $r_{H}$ vary from 0 up to 16.1 and 65.7 μmol L^-1^ d^-1^ over the 7-day incubation, respectively, and the turnover of the POC pool averaged 5.1 days.

### Turnover of the cell and viral fractions

Two pools are modeled here that track the ^13^C in the chemoautotrophs and the ^13^C in the viral particles, derived from Fig. 3 of the main text. Since the phage are labeled from the host’s DNA/RNA, that cellular pool gets labeled first, which then flows into the cell free viral particles, as in this diagram,

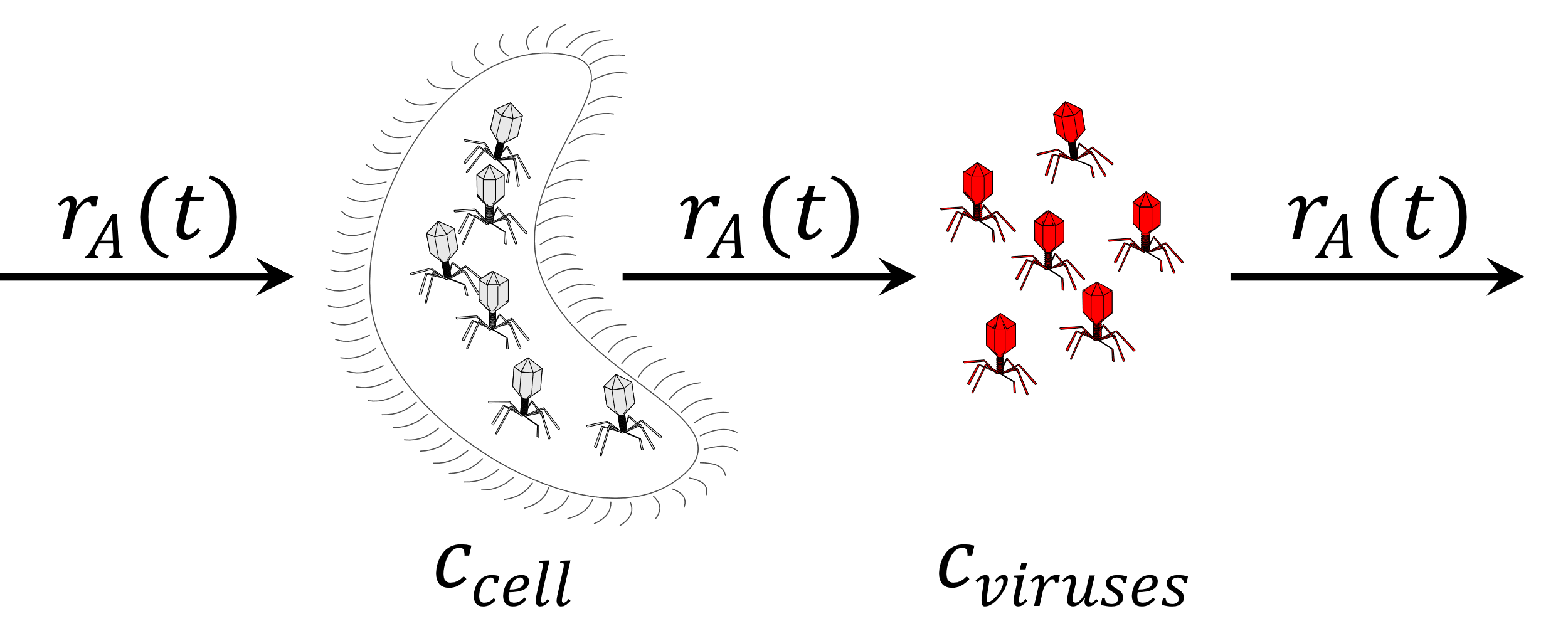

We will assume the pool concentrations (^12^C + ^13^C) are approximately at steady state, where the flow through the two pools is given by the chemoautotroph growth rate, $r_{A}(t)$, determined above. From Table S4, the cell-enriched ^13^C fraction, $f_{cell}(t_{f})$, attains a value of 0.155 and the virus-enriched fraction, $f_{vir}(t_{f})$, is 0.066. Both pools start at background ^13^C levers, $f_{C13}=0.011$, but the flow into $c_{Cell}$ is enriched to $f_{DI13C}=0.413$ at time zero. Two ODEs can be written, one for the concentration of ^13^C in the cells, ${{}^{13}c}_{cell}$, and one for the ^13^C in the free viruses, ${{}^{13}c}_{vir}$, as given by

$$\frac{d{{}^{13}c}_{cell}(t)}{dt}=\left( f_{DI13C}-\frac{{{}^{13}c}_{cell}}{c_{Cell}} \right)r_{A}\left( t \right); {{}^{13}c}_{cell}\left( 0 \right)=f_{C13}c_{cell}$$

(4)

$$\frac{d{{}^{13}c}_{vir}(t)}{dt}=\left( \frac{{{}^{13}c}_{cell}}{c_{Cell}}-\frac{{{}^{13}c}_{vir}}{c_{vir}} \right)r_{A}\left( t \right); {{}^{13}c}_{vir}\left( 0 \right)=f_{C13}c_{vir}$$

(5)

These ODEs can be solved analytically, but contain two unknowns, the total concentration of the chemoautotrophs, $c_{Cell}$, and the concentration of the viruses, $c_{vir}$; however, we know the ^13^C fraction in the cells and the viruses at the end of the incubation, so the two unknowns can be solved from these two equations,

$${{}^{13}c}_{cell}\left( t_{f} \right)=f_{cell}\left( t_{f} \right)c_{cell}$$

(6)

$${{}^{13}c}_{vir}\left( t_{f} \right)=f_{vir}\left( t_{f} \right)c_{vir}$$

(7)

Based on the determined concentrations of $c_{Cell}$ and $c_{vir}$ and the chemoautotroph rate, $r_{A}(t)$, the turnover of the bacteria and virus pools averaged over the 7 day incubations were estimated to be 15.7 and 7.2 days.
